# Repetition of rhythmic patterns fosters neural representation of musical meter

**DOI:** 10.1101/2024.12.09.627467

**Authors:** Emmanuel Coulon, Sacha Baum, Tomas Lenc, Rainer Polak, Sylvie Nozaradan

## Abstract

Music often entails perception of periodic pulses (hereafter meter) which serve as an internal temporal reference to coordinate movements to music. Crucially, meter perception arises even when the musical rhythm only weakly cues meter periodicities (i.e., syncopated rhythms). However, syncopated rhythms are often looped in music, suggesting that repetition of rhythmic patterns may facilitate meter perception by providing periodic cues at a slower, supra-second timescale. Here, we tested this hypothesis by recording separately electroencephalographic (EEG) and behavioral responses (finger tapping) while participants listened to different syncopated rhythmic sequences. These sequences either consisted of a repeated pattern (repetition of 4.8 and 9.6-s patterns) or were generated without repetition. EEG responses showed overall periodization of the rhythmic input, at periodicities corresponding to those expressed as the meter in behavioral responses, and in contrast with the weak cues to these periodicities in the rhythmic inputs. Most importantly, pattern repetition strengthened this neural representation of the meter, demonstrating that supra-second periodicities in the rhythmic input further enhance sub-second periodicities in neural activity. These findings thus highlight the multiscale nature of temporal processes at stake in processing musical rhythm, and, more generally, complex rhythmic inputs involved in interpersonal interaction and communication.

## 1. Introduction

Music has a profound and universal effect on humans. This effect often results from movement (e.g., instrument playing, hand-clapping, singing, etc.), and also motivates movements (e.g., dancing, head bobbing, foot tapping, co-performers instrument playing, etc.) that are coordinated with the rhythmic input^[1,2]^. For this rhythmic movement coordination to occur, an internal representation of time is required that is, to some degree, invariant to the specific temporal arrangement of musical events. This internal representation typically takes the form of meter, usually conceptualized as a set of temporally recurrent pulses. These pulses often form a nested scaffolding over multiple (sub- to supra-second) timescales, including a pulse layer standing as a particularly prominent time reference, usually called the beat^[3,4]^.

Notably, a representation of meter can arise in response to a wide variety of rhythmic stimuli, ranging from controlled isochronous sequences (e.g., metronomes) to spectro-temporally complex real-world performances (e.g., marching bands, large orchestras, etc.). Hence, the brain does not stop at building an internal representation of meter that corresponds one-to-one to periodic recurrences physically present in the rhythmic input, but rather goes beyond. This phenomenon is illustrated well with syncopated rhythms, which typically feature only weak acoustic cues to meter periodicities. Yet, syncopated rhythms are commonly used in music and do not prevent listeners from perceiving a metric structure, as embodied in the production of periodic movement along with these rhythms^[5–7]^.

Analogously, studies using electrophysiology have repeatedly demonstrated that brain activity elicited while participants listen to syncopated rhythms shows a selective enhancement of periodicities corresponding to the perceived meter. In other words, there is robust evidence that meter perception is tied to processes that periodize the rhythmic sensory input, thus shaping the neural representation towards a behaviorally-relevant internal template^[8–12]^. Together, these findings highlight the fact that internal representations of meter cannot be solely explained by lower-level sensory processing of prominent temporal cues of the input, but at least partially involve processes that enable a higher degree of invariance with respect to the input^[13]^.

Critically, meter perception concerns a specific temporal range lying approximately between 100 and 1800 ms (as delimited through ethnomusicological work, and behavioral experiments where participants were instructed to tap in synchrony along with isochronous rhythms^[14–18]^). Given that this temporal range covers a large span, from sub-second to supra-second scales, meter perception has been hypothesized to involve multiscale temporal scaffolding processes whereby the temporal structure of the rhythmic input over this whole temporal range, from slow to fast, would serve as temporal cues to map metric pulses^[4,19–21]^.

Here, we aimed to move a critical step forward in investigating this multiscale temporal scaffolding with a focus on its slower, supra-second end. We first propose a scheme where this temporal range for meter is organized into three temporal timescales or layers, from slow to fast (Fig. 1). This scheme then serves as a basis to manipulate the prominence of the temporal cues at the slow supra-second layer while keeping parameters at the faster sub-second layers constant, thus informing on the contribution of the supra-second layer to the multiscale temporal scaffolding process at stake in meter perception.

**Figure 1:**
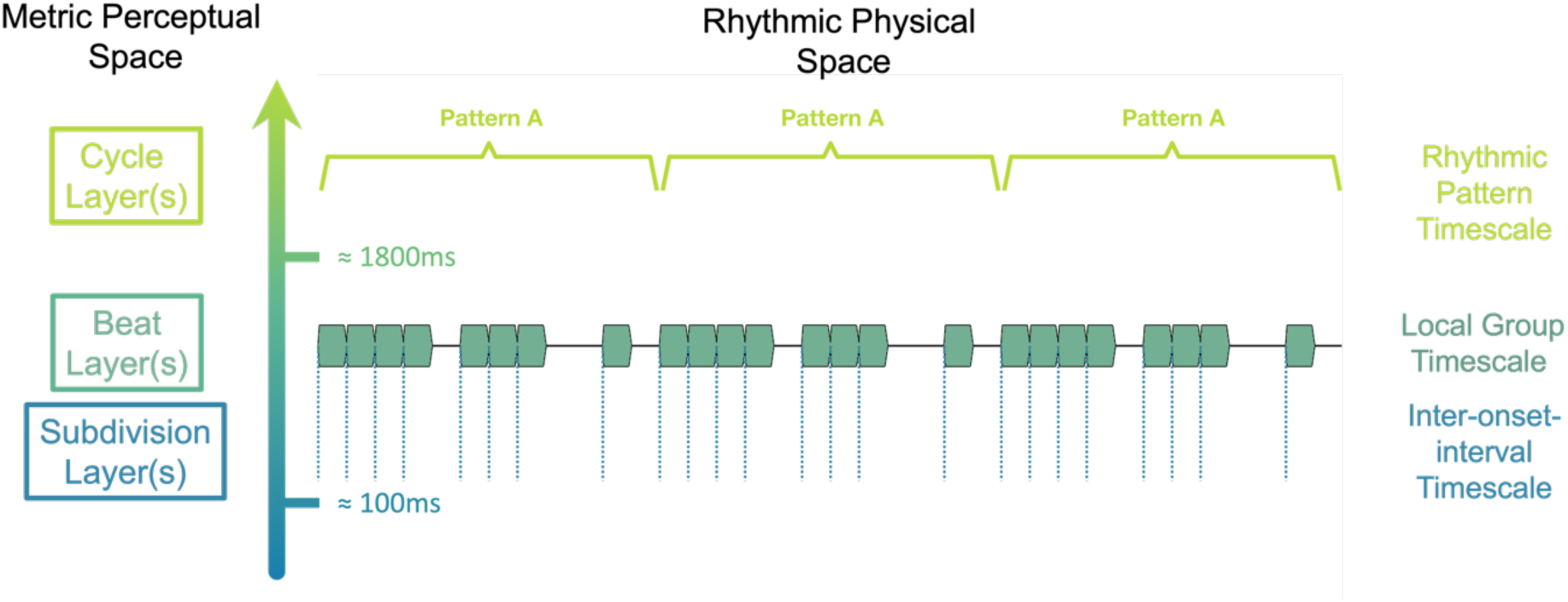
Three-layer scheme of the temporal range for meter processing, showing the physical space of the rhythmic input and corresponding metric perceptual space. The figure depicts an example rhythmic sequence with prominent periodic cues in the faster layer, formed by recurrent intervals between successive sound events (dashed vertical lines) and prominent periodic cues in the slower layer, formed by the seamless repetition of the rhythmic pattern (as depicted with squared brackets). In contrast, the intermediate layer does not show prominent periodic cues due to the local arrangement of sound events forming a syncopated rhythm.

As depicted in Figure 1, the proposed scheme comprises a faster timescale (hereafter referred to as *subdivision layer*) concerned with single time intervals delimited by the successive events making up the rhythmic input, which can serve as a basis to map fast metric pulses^[4,16,17]^. The intermediate timescale (hereafter called *beat layer*) is concerned with the local arrangement of these successive events (also referred to as groups)^[4]^. These local arrangements can be considered as unsyncopated or syncopated (also referred to as strongly periodic or weakly periodic in the beat layer), depending on the extent to which these groups match the periodicity of the perceived beat^[22,23]^. Finally, the slower timescale (hereafter referred to as *cycle layer*) is concerned with longer, seamlessly repeating, or looped, rhythmic patterns, which can serve as a basis to map slow metric pulses^[4,15,24]^. Notably, the boundaries between these three timescales certainly depend on a multitude of factors pertaining to the stimulus (e.g., speed, timbre, etc.) and to the listener (e.g., cross-cultural or inter-individual differences, knowledge of the perceptual conventions inherent in musical genres, etc.). Nevertheless, there is generally a considerable overlap between the range of temporal cues of the rhythmic input and the range of perceived metric pulses, particularly at the beat and subdivision layers^[2,4,12]^.

Thus far, most neuroimaging studies focused on the role of temporal cues at the intermediate timescale corresponding to the beat layer, while keeping periodic recurrence at the slower timescale highly prominent through the use of looped rhythmic patterns^[8,10,11,13,25,26]^. However, little effort has been focused on evaluating the effect of supra-second temporal cues on the internal representation of meter. Yet, repetition of rhythmic patterns represents a core component of music^[24,27–29]^, one that is key to the aesthetic appeal and social power of music^[5,30]^. Repeating rhythmic patterns are often easily recognized and learned without requiring explicit training, and this periodic recurrence can help to structure the musical piece and anticipate future events^[4,27,31]^.

Therefore, we hypothesize that the periodicity provided by the repetition of rhythmic patterns serves as temporal cues to the meter. These cues would offer a temporal anchor point onto which faster metric pulses at the beat and/or subdivision layers can be mapped by interpolation, through multiscale temporal scaffolding potentially involved in meter perception^[4,32,33]^. We tested the role of these supra-second temporal cues by recording behavioral and neural responses in human healthy adult volunteers. Behavioral measures of the internal meter representation were obtained by asking participants to tap the finger along with the meter they would perceive while listening to rhythmic sequences. In a separate session, neural responses were captured using electroencephalography (EEG) while the same participants were listening to the same rhythmic sequences, from which neural representation of the perceived meter were measured using frequency analysis (see^[12,34]^ for reviews).

These behavioral and neural measures were compared across three sequences presented in separate conditions. In the three sequences, temporal cues to the beat layer were invariantly low (i.e. weakly periodic, syncopated rhythms), whereas the periodic recurrence in the slower timescale corresponding to rhythmic patterns was manipulated such as to gradually decrease across stimuli (sequence #1: repetition of a 4.8 s pattern, sequence #2: repetition of a 9.6 s pattern, and sequence #3: no pattern repetition). A decrease of the behavioral and neural measures of internal meter representation across conditions would thus demonstrate a critical role of slow periodic cues in meter processing. Alternatively, observing comparable meter-related responses across conditions would suggest that periodic cues at the faster beat and/or subdivision layers may be sufficient to enable meter representation.

We also tested how these supra-second temporal cues interacted with other factors assumed to play a role in meter processing, namely short-term prior context^[35]^ and long-term musical expertise. To this aim, different presentation orders of the conditions were compared, thus allowing to test whether supra-second temporal cues would be used as temporal anchor for meter not only when these slow cues are actually present in the sequences but also in subsequent sequences lacking these cues (Fig. 3). Additionally, group comparison across professional musicians and non-musicians aimed to test the extent to which long-term musical practice overrides the lack of slow temporal cues in the rhythmic input, allowing for the representation of a stable internal metric template irrespective of sensory input degradation^[2,36]^.

## 2. Materials and Methods

### Participants

To measure long-term shaping of meter perception by music practice, two groups of 26 healthy volunteers recruited from Brussels area (Belgium) participated in this study. The first group consisted of professional musicians (17 females, 5 left-handed, 27.54 ± 6.18 years old, 19.5 ± 5.46 years of musical practice, and 26.75 ± 7.8 hours of musical practice per week - mean ± standard deviation) who were either studying or had graduated from a music conservatory (i.e., bachelor’s or master’s degree) at the time of the experiment. The second group consisted of non-musicians (17 females, 2 left-handed, 25.96 ± 5.53 years old, 0 ± 0 years of musical practice, and 0 ± 0 hours of musical practice per week). Both groups were matched for dance experience (Musicians: 2.34 ± 3.89 years of dance experience and 1.15 ± 2.96 hours of weekly practice; Non-musicians: 0.81 ± 2.00 years of dance experience and 0.60 ± 1.16 hours of weekly practice; both p-values > 0.05). Participants had no history of hearing, neurological or psychiatric disorders, and gave a written informed consent before the start of the experiment. This study was approved by the local ethics committee (ref. 2018-353) and all experiments were performed in accordance with relevant guidelines and regulations.

### Auditory Stimuli

A set of 14 seed rhythms was created using Matlab (R2020a, *The MathWorks, Natick, USA*; Fig. 2). Each rhythm was based on a 12-interval grid structure made of 200 ms evenly spaced time points and consisted of eight sound events, and four silent events. For each seed rhythm, the arrangement of sound and silent events on the 12-interval grid was manipulated to obtain 2.4 s syncopated (i.e., weakly periodic in the beat layer) rhythms.

**Figure 2:**
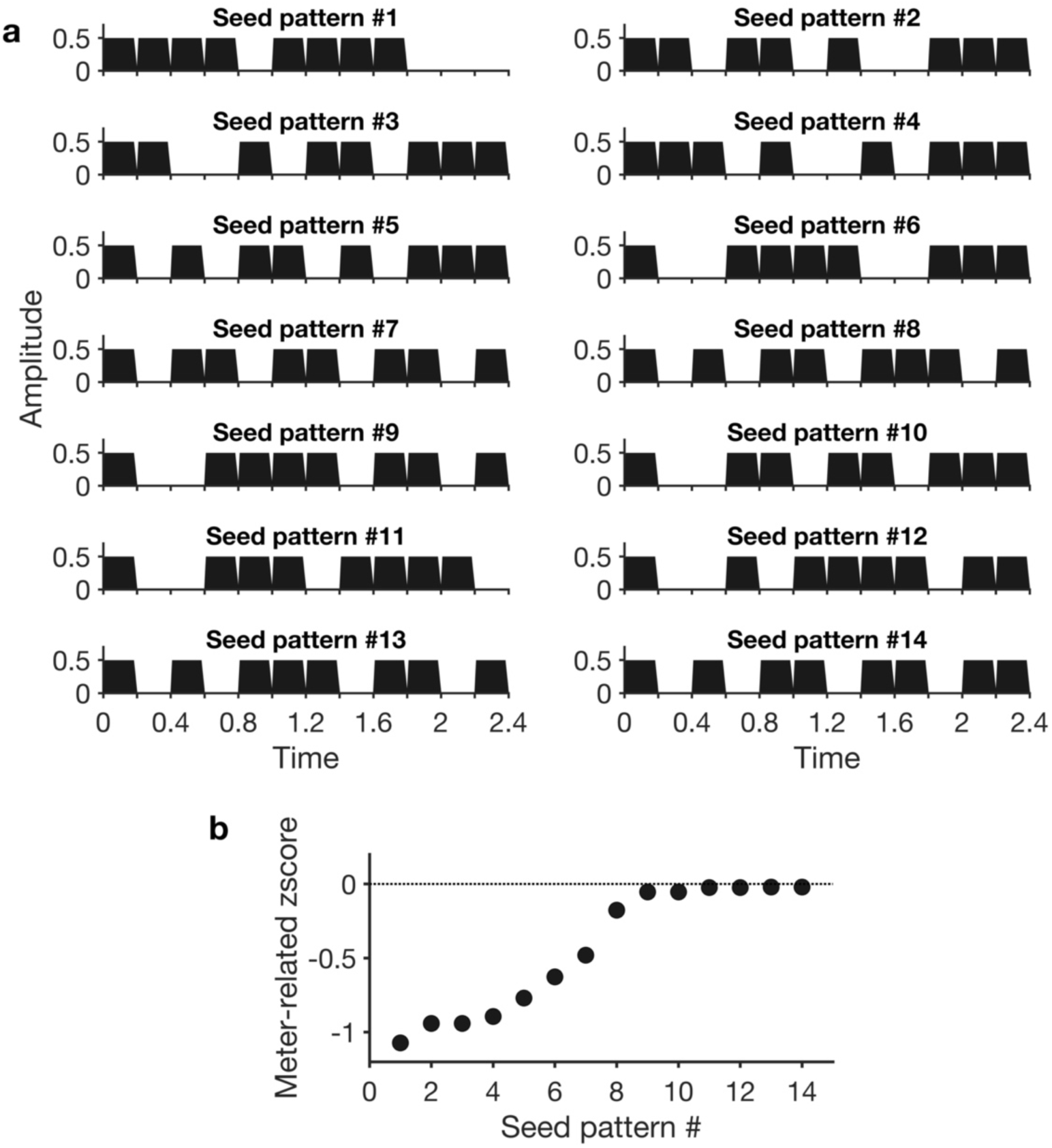
Temporal envelope of the fourteen seed rhythms. (a) All rhythms were based on a twelve-interval 200 ms grid structure (i.e., 2.4 s in total), with eight of the intervals consisting in sound events and the remaining four intervals consisting in silent events. The sound and silent intervals were arranged such as to yield syncopated rhythms. (b) Weak prominence of beat periodicities was confirmed through frequency analysis of the amplitude modulation of the rhythms, from which meter z-scores were calculated All seed rhythms showed negative meter-related zscores, indicating weak prominence of meter-related periodicities in their modulation spectrum compared to meter-unrelated periodicities.

Sound events consisted in a pure tone of 200 ms duration (10 ms and 20 ms linear ramps up and down). The carrier frequency of the pure tone varied between 150 and 200 Hz ([150.00, 161.19, 173.21, 186.12, 200] Hz) in each successive trial to increase participants’ vigilance due to the change in tone between trials. These relatively low tone frequencies were chosen since low tones have been shown to elicit brain responses with higher signal-to-noise ratios^[37]^, and stronger meter-related neural responses^[8]^. To account for the spectral sensitivity of the auditory system^[38]^, the intensity of each pure tone was equalized to 70dB_A_ using a Bruel & Kjaer (type 2250, *Denmark*) sound level meter.

To confirm that temporal cues to beat-layer periodicities were weak in these stimuli (i.e., syncopation criterion), we measured the prominence of meter periodicities in the modulation spectrum of each rhythm. This was done by extracting the envelope of the acoustic signal of each 2.4 s rhythmic pattern using a Hilbert transform and subsequently applying a Fast Fourier Transform (FFT). The distribution of energy within the first 12 frequency bins (from the frequency corresponding to the duration of the rhythmic pattern, here 1/2.4 s = 0.41 Hz, to the 11th harmonics corresponding to the period of the grid timepoints, 1/0.2 s = 5 Hz) reflects periodic recurrences in the amplitude modulations of the acoustic signal^[9,34,39,40]^.

From this set of frequencies of interest, frequencies were classified as meter-related when corresponding to the most plausible metric pulses (1.25Hz and harmonics, based on tapping behavior observed in studies using similar rhythms^[8,10,13]^), and as meter-unrelated frequencies otherwise. To measure the relative prominence of meter-related frequencies, amplitudes at all frequencies of interest were converted to z-scores and the z-scores at meter-related frequencies were averaged to serve as an index of their relative prominence in the modulation spectra (see^[34]^ for further details on this approach). All seed rhythms showed negative meter-related z-scores, indicating that the amplitude at meter-related frequencies was, on average, smaller than the average amplitude across all frequencies of interest. In other words, meter-related frequencies did not stand out relative to the whole set of frequencies characterizing the temporal structure of the rhythmic patterns, thus confirming the syncopation criterion.

The 14 seed rhythms were then concatenated in three 67.2s sequences that varied in terms of pattern repetition. The “medium pattern repetition” sequence consisted of a 4.8-s rhythmic pattern made of the concatenation of two of the seed rhythms (seed rhythms #5 and 10), which was seamlessly looped 14 times within the sequence. The “long pattern repetition” sequence comprised the same 4.8s pattern from the medium pattern repetition sequence followed by two additional seed rhythms, thus forming a longer rhythmic pattern of 9.6s (seed rhythms #5, 10, 8, 9) which was seamlessly looped 7 times within the sequence. Lastly, the “no pattern repetition” sequence was made by randomly concatenating the 14 seed rhythms to generate the 67.2s sequence (seed rhythms #6, 8, 11, 5, 12, 5, 7, 6, 11, 7 etc.).

All sequences were controlled (i) to yield similar meter-related z-scores, (ii) to not present identical rhythms following each other within a pattern, and (iii) so that the no-pattern repetition sequence would not reproduce parts of the pattern from the other two sequences. Note that the label “short pattern repetition” sequence was purposely avoided here, as it better refers to the duration of the seed rhythms (i.e., 2.4-s), which corresponds to the typical duration of rhythms used in previous similar neuroimaging studies^[8,10,13,41]^. Note also that a condition with repetition of such a 2.4 s rhythmic pattern was not included in the current study to maintain a reasonable experiment duration, and because longer patterns were hypothesized to more effectively target the 6-8 s limit for the implicit learning of rhythmic patterns^[15,42]^. Audio files of the stimuli are available on an online repository (https://zenodo.org/records/13870035?preview=1).

To test whether low-level sensory processing from early stages of the auditory pathway could explain substantial enhancement of meter frequencies as compared to the input^[43]^, all three sequences were analyzed using a cochlear model to estimate the sound representation at the level of auditory nerve fibers (as implemented in the Auditory Toolbox for Matlab, with 128 characteristic frequencies distributed from 130 to 16.000Hz^[44]^). The output of this model was averaged across the cochlear channels, and the signal was transformed into the frequency domain by applying an FFT. The meter-related z-score was then computed for each stimulus condition to confirm the lack of prominent meter-related frequencies, thus excluding possible low-level confounds in the EEG results.

### Experimental design

The experiment took place in a single session. Each participant listened to all three sequences presented in blocks, and the block corresponding to the “long pattern repetition” condition was presented twice (Fig. 3) to test for effects of short-term context. Another way to test for possible effects of short-term context was to compare two different fixed orders of the blocks, with the “no pattern repetition” condition placed either at the beginning (i.e., no prior context), or at the end of the experiment (i.e., maximum prior context). Within each group of participants (musicians vs. non-musicians), half of each group was assigned to a different condition order, thus yielding a mixed factorial design.

**Figure 3:**
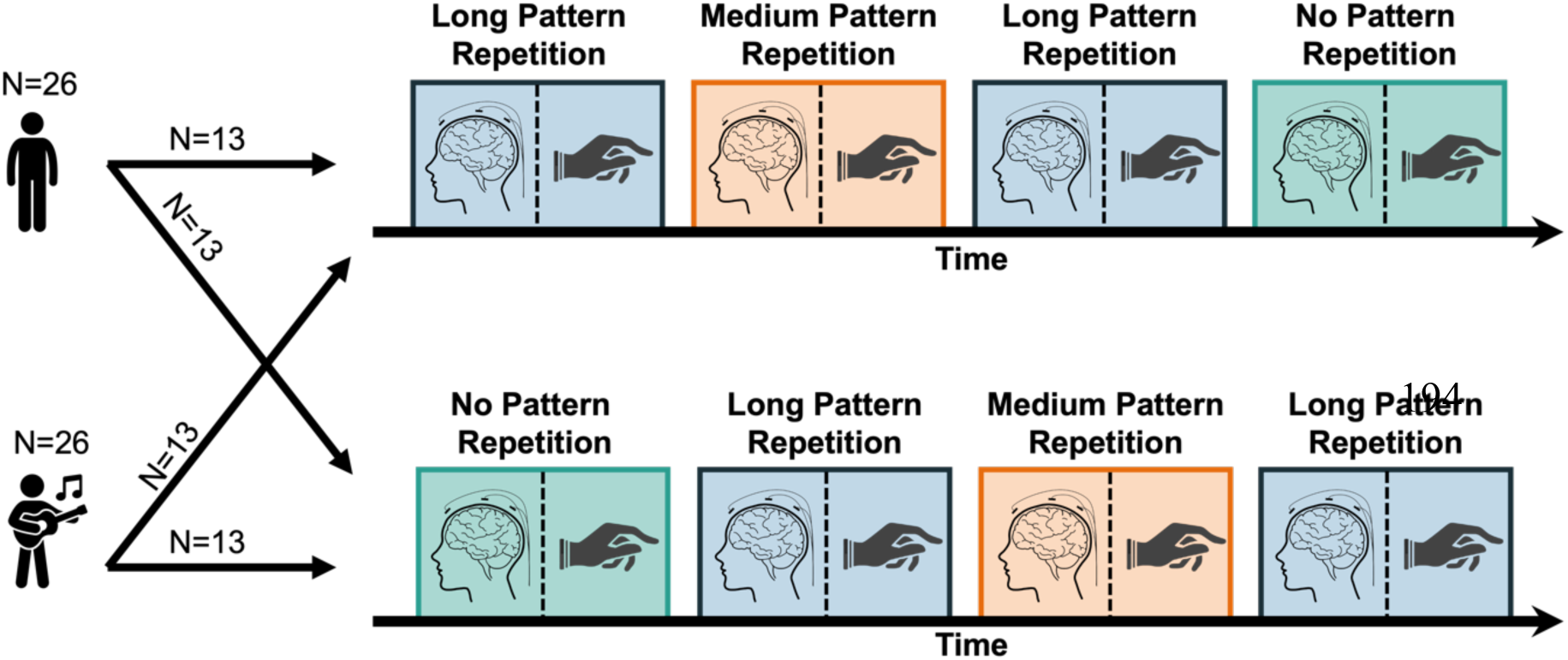
Experimental design. Musician and non-musician groups of participants were presented with four blocks in two possible fixed orders - the top and bottom condition orders providing maximum and minimum prior context respectively, with the “no pattern repetition” condition positioned either at the end or the beginning of the block design. Half of each group was assigned to each condition order, and for each condition neural activity was recorded using electroencephalography, followed by a tapping session.

Each of the four resulting blocks consisted of 10 EEG trials followed by 3 tapping trials. During EEG trials, participants were seated in a comfortable chair and were asked to listen carefully to the rhythmic sequences while fixating a visual target placed in front of them and avoiding any unnecessary head or body movements. To maintain participants’ attention to the temporal properties of the stimuli during EEG trials, participants were asked to report whether they heard a tempo change, and if so the direction of this change, at the end of each trial^[8,35,44]^. These tempo changes were introduced in an additional sequence snippet appended to the end of 6 out of 10 trials (3 accelerations, 3 decelerations, and 4 trials without tempo change). Modulations of the tempo were implemented following a 4.8-s long cosine waveform with a fixed magnitude for musicians (minimum and maximum grid intervals of 188.7 and 217.4 ms respectively), and an adaptive two-down, one-up staircase method^[45]^ (grid intervals starting at 169.5 for accelerations and 238.1 ms for decelerations with adaptative steps of 8ms) for non-musicians to account for their reduced ability to detect tempo modulations as compared to musicians^[46]^. Note that these additional snippets were excluded from further analyses, thus exclusively leaving responses to rhythmic input with a steady tempo for the analyses. Before the start of the experiment, participants were familiarized with several examples of these tempo changes applied to syncopated rhythms that were not used in the actual experiment.

For the tapping trials, participants were instructed to tap the regular pulse that was spontaneously perceived while listening to the stimulus by producing up and down movements with the index finger of their preferred hand on a custom-built response box (i.e., the exact instructions were: “*Tap the index finger of your preferred hand in time with the regular pulse that you perceive when listening to the rhythm, as if you were clapping your hands on the music during a live performance*”). The experimenter remained in the recording room with the participant at all times to monitor compliance with the procedure and instructions.

### Tapping Recording and Analysis

Tapping responses were recorded using a custom-built response box made of a hard tapping surface (i.e. producing auditory feedback, mitigated by the ear inserts used to deliver acoustic inputs simultaneously) and a pressure sensor underneath. From the tap time series, intertap intervals (ITIs) were calculated as durations between successive taps. For each trial, participant and condition, the median ITI was taken as an index of the perceived beat period^[2]^. Additionally, we converted the tap time series to a sequence of phases relative to the predicted beat-layer pulse frequency, determined by the meter periodicity closest to each participant’s median ITI. The circular mean was then taken as an index of tapping stability, with values ranging from 0 to 1, respectively, indicating low and high synchronization with the rhythmic input^[7,47]^.

### EEG Recording and Preprocessing

The EEG was recorded using a Biosemi Active-Two system (Biosemi, Amsterdam, Netherlands) with 64 Ag-AgCl electrodes placed on the scalp according to the international 10/20 system, and two additional electrodes placed on the mastoids. EEG recordings were preprocessed using Matlab (R2020a, The MathWorks, Natick, USA) and Letswave (http://letswave.org).

First, signals were digitized at a sampling rate of 512 Hz. To remove slow drifts, EEG recordings were high-pass filtered offline at 0.1 Hz (4th order Butterworth filter), segmented from 0 to 67.2 s relative to the trial onset (excluding any appended snippets with tempo changes relevant for the perceptual task) and re-referenced to the average of mastoid electrodes to increase the signal-to-noise ratio of neural responses to acoustic rhythms in frontocentral channels, which would be later selected^[48,49]^.

Excessively noisy channels were visually selected and linearly interpolated across all trials and conditions, separately for each participant (maximum 3 channels interpolated in 2 participants). Artifacts attributed to eye blinks and horizontal eye movements were identified and removed using independent component analysis^[50]^ based on visual inspection of their typical waveform shape and topographic distribution. For each participant and condition separately, the epochs were averaged across trials in the time-domain to reduce the contribution of background noise that was not time-locked to the stimuli, thus improving the signal-to-noise ratio of the EEG activity^[51]^.

The signal was then averaged across a set of 9 frontocentral channels (‘F1’, ‘Fz’, ‘F2’, ‘FC1’, ‘FCz’, ‘FC2’, ‘C1’, ‘Cz’, ‘C2’ in the 10-20 international system) based on prior observations of brain responses to rhythmic sound patterns being predominantly captured by these channels^[8,10,35]^. Finally, the averaged waveforms were transformed into the frequency domain using an FFT, and baseline corrected by subtracting the average amplitude at the 2nd to the 5th neighboring bin on both sides relative to each frequency bin to locally correct for the background noise in the EEG signal^[51]^.

### Relative amplitude of the meter-related neural activity in the frequency domain

A vast majority of previous EEG studies using a frequency-tagging approach to investigate the neural representation of meter have used repeated rhythmic patterns^[8,10,11,13,25,26]^. In such a context, the selection of frequencies of interest is relatively straightforward, and directly based on the fact that the response to a repeated rhythmic pattern is elicited repeatedly with a fixed a priori known period. Hence, any response reliably elicited by the pattern should itself repeat at the rate of pattern repetition in the sensory input. Such a systematically repeated response can be conveniently isolated in the frequency domain, since it only contains peaks at frequencies corresponding to the pattern repetition rate and harmonics.

Critically, the distribution of amplitudes across these frequencies of interest is related to the shape of the response elicited *within* each repetition of the rhythmic pattern. Moreover, the degree of periodic recurrence of the response (i.e., meter periodicities) within each pattern cycle is reflected in the relative prominence of amplitudes at a subset of frequencies of interest (i.e., meter-related frequencies), which correspond to the rate of this nested recurrence and its harmonics. To measure such prominence at the rate of the perceived metric periodicities (as directly informed by the group-averaged median ITI), the amplitudes across all frequencies of interest are typically standardized as z-scores and averaged across meter-related frequencies (i.e. frequencies corresponding to the rates and harmonics of the metric pulses). Notably, the resulting mean z-score quantifies the amplitude at meter-related frequencies *relative to* the amplitude at other, meter-unrelated frequencies, which are expected to emerge in the response spectrum due to stimulus repetition, but do not capture meter periodicities. The mean meter-related z-score thus constitutes a normalized index of the prominence of meter periodicities in the EEG signal^[9,12,34,52]^.

Along this line, we determined a set of meter-related frequencies based on the tapping, namely the observed convergent ITIs across conditions. However, determining meter-unrelated frequencies based on the harmonics of the pattern repetition rate would be ill-suited here, as each condition contained a different repetition rate (up to a complete absence of pattern repetition), thus precluding comparisons across conditions due to the profound differences in the number of meter-unrelated frequencies. Yet, meter-unrelated frequencies are key to standardizing response amplitudes at meter-related frequencies, thus capturing the prominence of meter periodicities irrespective of unit and scale^[34]^. Therefore, we adopted a new approach to selecting frequencies of interest and overcoming this imbalance without any prior assumption about possible meter-unrelated frequencies.

First, we identified the most prominent peaks from the group-averaged EEG spectrum of each condition. These were obtained by running the *findpeak* function implemented in Matlab, with a minimum peak prominence set as the mean plus two standard deviations of the spectrum between 0 and 30Hz. This measure was computed separately for each group of participants, and only frequencies present in both groups were retained. The same procedure was applied to the modulation spectrum of each of the three rhythmic sequences, and the series of prominent peaks from the three stimuli were merged with those extracted from their corresponding EEG spectrum to better capture the input/output transformation. This way, small peaks at the stimulus level that are boosted in the EEG, or otherwise, prominent peaks in the input that are reduced in the output are both considered as frequencies of interest.

Furthermore, peaks obtained below 1.25 Hz (i.e., pulse period corresponding to 4 events, or 800ms) were excluded from further analysis since they were expected to be particularly affected by 1/f noise in the EEG signal. In addition, peaks at 5Hz and harmonics were also excluded as they are likely to be mostly driven by the shape of the average response to individual sound events making up the stimuli.

The steps describe thus far could, in principle, yield a rather different number of frequencies of interest across conditions. To avoid such imbalance, we retained only the smallest number of frequencies of interest among conditions by (i) selecting those with the largest amplitude in the modulation spectrum of each stimulus, and (ii) forcing the inclusion of meter-related frequencies corresponding to median ITIs tapped by participants (i.e., 1.25 and 2.5 Hz), resulting in 3 meter-related and 24 meter-unrelated frequencies.

Finally, the amplitudes of all these frequencies of interest were converted into z-scores and averaged separately for the meter-related frequencies (frequencies corresponding to the period of the median ITIs, and harmonics – 1.25, 2.5 and 3.75 Hz) and meter-unrelated frequencies (all non-meter-related frequencies of interest). As expected, this procedure yielded a selection of different meter-unrelated frequencies across conditions. However, the fact that this selection was conducted within an identical spectral range and was restricted to an identical number of these frequencies across conditions ensured fair comparison of the relative prominence of meter-related frequencies across conditions.

### Statistical analysis

Statistical analyses were performed using R (version 4.2.1) with the significance level set at p < 0.05. To account for the absence of normal distribution in behavioral results, non-parametric Mann-Whitney tests were used for comparisons across the two groups, and between the two condition orders, and a non-parametric Friedman test was used to compare across conditions. For neural responses, a three-way mixed ANOVA was conducted to evaluate the main effect and interaction of the factors condition, musical expertise, and condition order. Where relevant, the Greenhouse–Geisser correction was used to correct for violations of sphericity in the performed ANOVAs. Post-hoc comparisons were carried out using paired t-tests, and a false discovery rate (FDR) correction was applied when needed to adjust for multiple comparisons.

## Results

### Behavioral results

Figure 4A depicts the distribution of inter-tap intervals (ITIs) for both groups of participants across all four conditions. The observed convergent median ITIs at the group level and across conditions were thus used to inform the selection of meter-related frequencies, corresponding here to 1.25 Hz (i.e., 1/800 ms, or 4 times the grid interval) and harmonics (thus including also 2.5 Hz, corresponding to 1/400 ms, or 2 times the grid interval) for the frequency analysis of neural responses, in line with previous studies^[13,35]^.

**Figure 4:**
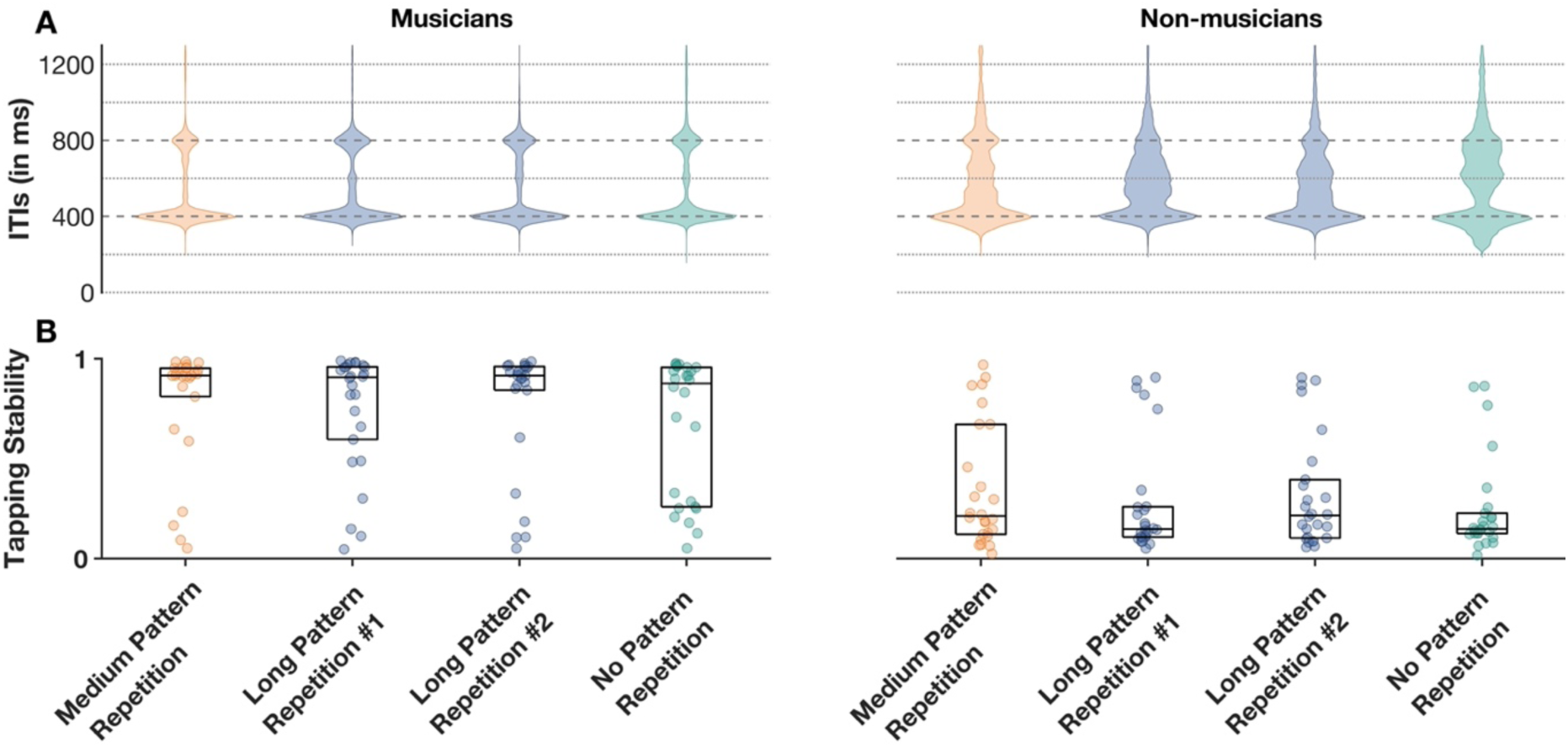
Behavioral results: non-musicians showed overall poorer stability at tapping the beat along with the rhythmic inputs, as compared to musicians. (A) Distribution of intertap intervals (ITIs) across participant groups and conditions. Horizontal lines correspond to integer multiples of the 200-ms grid structure on which rhythmic patterns were built, thus corresponding to plausible beat rates participants could have synchronized their tapping to. Dashed lines highlight 400- and 800-ms, corresponding to the convergent tapping rates across participants. (B) Tapping stability. Values ranging from 0 to 1 indicate low and high synchronization with a steady periodic beat, respectively. The horizontal line and limits of the boxplot indicate the median and 25^th^ and 75^th^ percentiles respectively.

To evaluate behavioral responses, the tapping stability (i.e., vector strength of the circular asynchrony between tapping onsets and the nearest hypothetical pulse position) was compared between groups, condition orders, and across conditions (Fig. 4B). Professional musicians showed higher stability than non-musicians in all conditions (Medium Pattern Repetition: W = 564, p = 1.442e-05, 17^2^ = 0.574 ; Long Pattern Repetition #1: W = 578, p = 3.304e-06, η^2^ = 0.609; Long Pattern Repetition #2: W = 574, p = 5.111e-06, 17^2^ = 0.599; No Pattern Repetition: W = 580, p = 2.644e-06, 17^2^ = 0.614). However, within-group comparisons across conditions, and condition orders showed no significant effect (Condition, Musicians: χ^2^(3) = 1.57, p = 0.67; Condition, Non-musicians: χ^2^(3) = 3, p = 0.39; Condition Order: W = 5784, p = 0.387), though some non-musicians appeared to strongly benefit from the order offering maximum prior context, leading to a bimodal distribution of tapping asynchronies and a close to significant interaction (Order, Non-musicians: W = 1608, p = 0.09).

Together, these results suggest that degradation of pattern repetition, as well as prior context, did not play a significant role in tapping performance. Only musical expertise had a positive effect, as observed by the more clustered ITIs and increased tapping stability measured in the musicians group.

### Frequency Analysis of the Stimulus and EEG responses

Figure 5 displays the modulation spectra across stimulus conditions obtained from the cochlear model (top row) and EEG responses (musicians and non-musicians in the middle and bottom rows respectively). As expected, the degradation of pattern recurrence causes the acoustic energy (and the corresponding output of the cochlear model) to be spread across a larger number of frequency bins that is directly determined by the duration of the repeated pattern in the sequence (Fig. 5, top row), hence the necessity for a proper adjustment of a method for selecting frequencies of interest as described in the materials and methods section. After running the findpeak function on the modulation spectrum of each stimulus condition and their corresponding group-averaged EEG spectra, the medium pattern repetition condition showed the fewest prominent peaks (n=27), which determined the number of frequencies of interest that would then be selected in each condition. After classifying 1.25Hz and harmonics as meter-related frequencies (Fig. 5 in red, n=3, as informed by tapping results), and all other frequencies of interest as meter-unrelated (Fig.5 in blue, n=24), the modulation spectrum of the cochlear model responses showed a weaker prominence of meter-related frequencies compared to meter-unrelated frequencies. This observation was confirmed by the negative meter-related z-score obtained in all four conditions (Fig. 6, horizontal black line), indicating that the rhythmic sequences contained little cues to the metric pulses as tapped by most participants.

**Figure 5:**
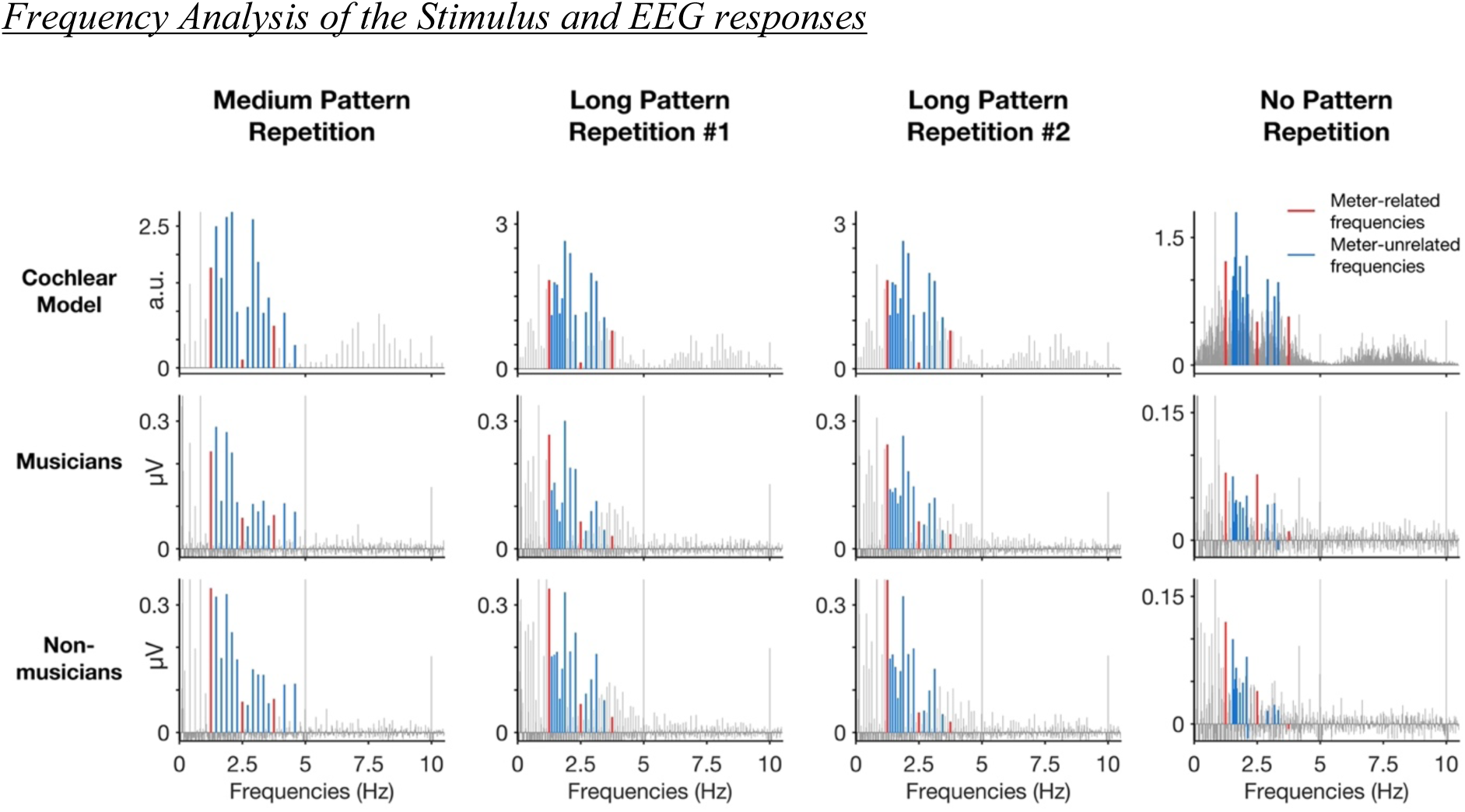
Modulation spectra obtained using cochlear model for each stimulus condition (top row), and their corresponding EEG responses in musicians (middle row), and non-musicians (bottom row). Frequencies of interest are respectively highlighted in red or blue for meter-related or meter-unrelated frequencies. EEG spectra correspond to a pool of 9 frontocentral channels and are averages over the 26 participants in each group.

**Figure 6:**
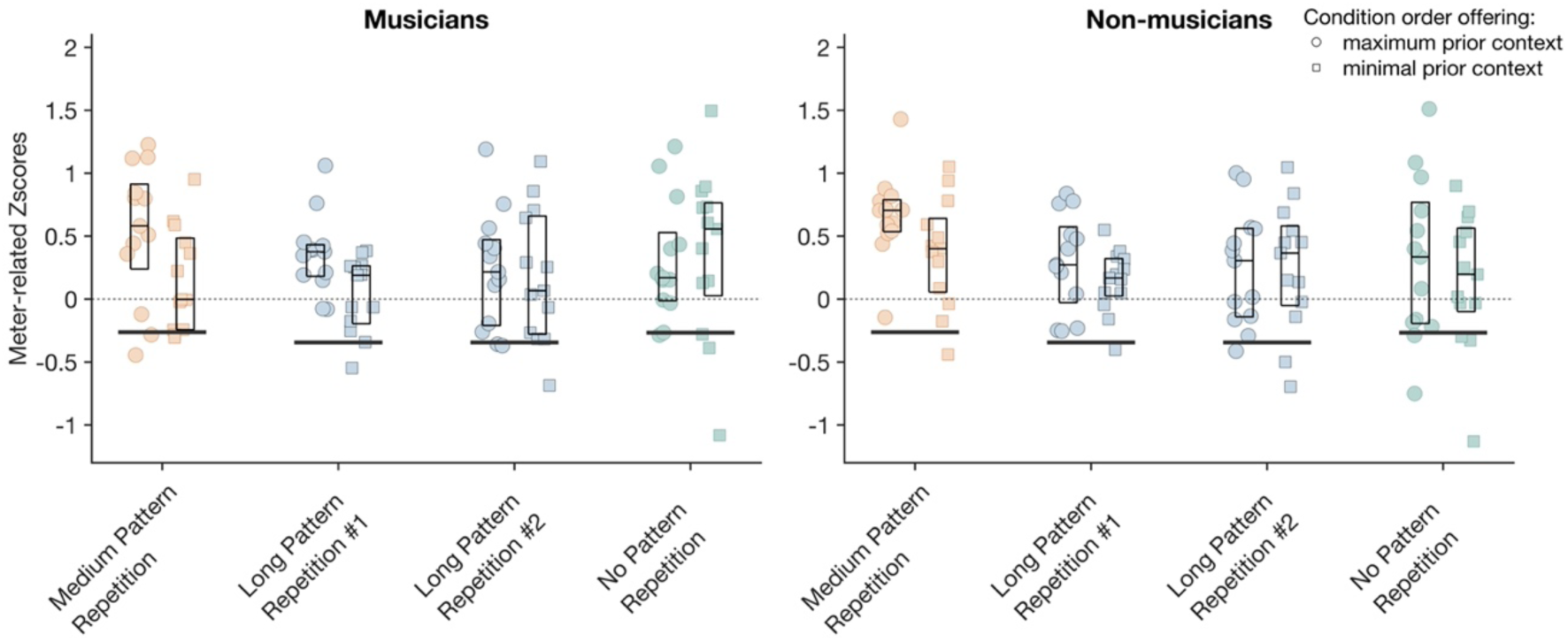
Mean z-score of EEG responses at meter-related frequencies across groups (left plot: musicians, right plot: non-musicians), for all conditions and condition order (circles: condition order maximizing prior context, squares: condition order minimizing prior context). Corresponding values obtained from the cochlear model output are shown as vertical black lines.

One-sample one-tailed t-tests were then performed to estimate whether meter-related frequencies were more prominent than predicted by the cochlear model, which could not be explained by lower-level tracking of prominent acoustic features of the stimuli^[12]^ (Fig. 6, black horizontal line). Both groups showed a selective enhancement of meter-related frequencies in the EEG in all four conditions and for both condition orders (all p-values < 0.05, FDR corrected, Table 1). These results align with previous work showing a periodization of the input, even when the input shows weak meter periodicities in its modulation spectrum^[9,11,12]^. However, these results go one step further by showing that the processes underlying periodization persist in response to stimuli with degraded periodicities at slower timescales, where periodic recurrences associated with pattern repetitions typically occur.

**Table 1:**
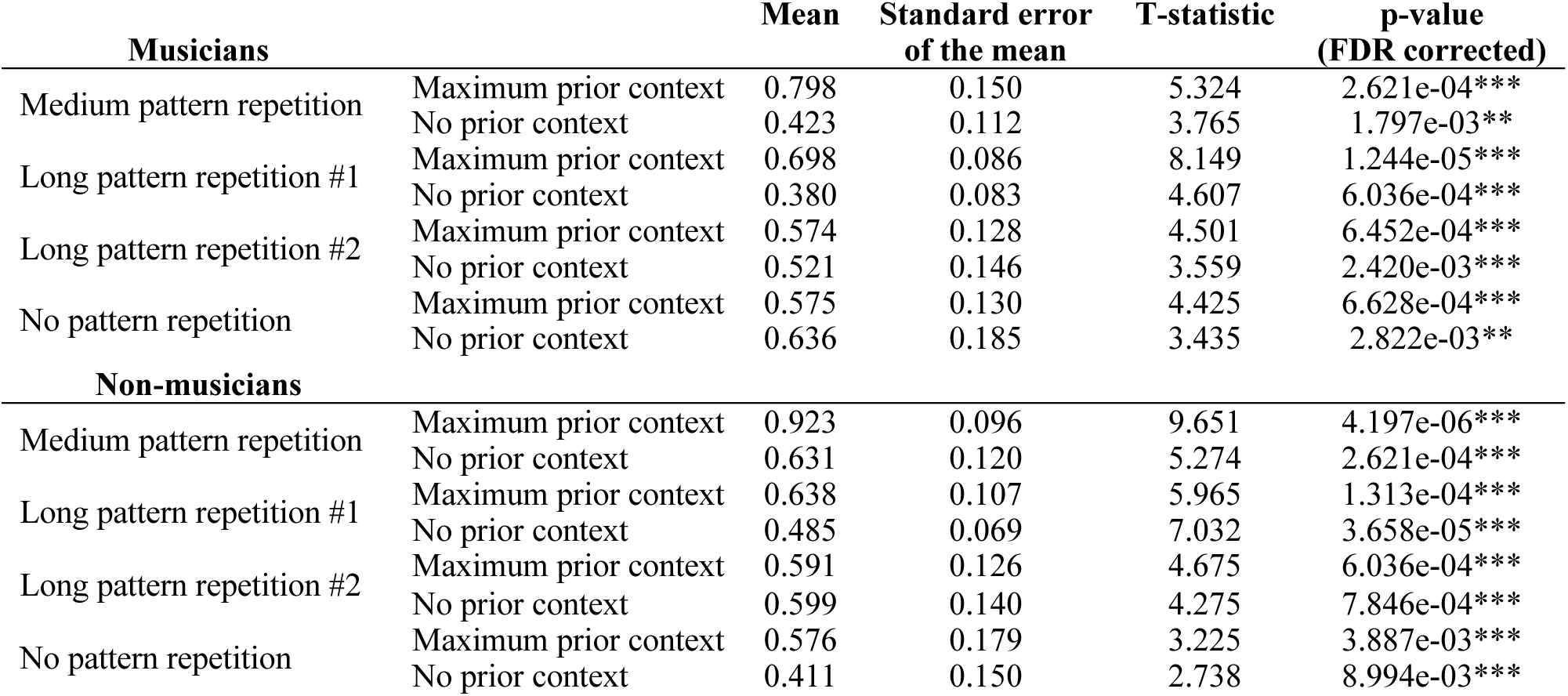
Selective enhancement of EEG meter-related zscores observed in each group, condition and condition order. The p-value is flagged with one star (*) if lower than 0.05, two stars (**) if lower than 0.01, and three stars (***) if lower than 0.001.

Finally, and more central to the aim of the current study, we estimated the effect of these slow temporal recurrences, along with effects of musical expertise and prior context, on the amplitude of EEG responses at meter-related frequencies (Fig. 6). The three-way mixed ANOVA with the factors condition, group, and condition order showed a main effect of condition (F(3,48) = 2.711, p = 0.0470), with larger meter-related z-scores observed in the medium-pattern repetition compared to the first presentation of the long-pattern repetition condition (t(51) = 3.411, p = 0.006, 17^2^ = 0.472), as well as a main effect of condition order (F(1,48) = 4.828, p = 0.033), with greater enhancement of meter-related frequencies in the order that provided maximum prior context (i.e. starting with the medium pattern repetition condition; t(103) = 2.553, p = 0.006, 17^2^ = 0.250). However, no main effect or interaction was found for the group factor, suggesting that listeners’ neural responses may be similar irrespective of levels of musical expertise.

## Discussion

The current study investigated the role of rhythmic pattern repetition in meter processing. As hypothesized, pattern repetition was found to strengthen neural representation of the meter. Moreover, we observed stronger neural emphasis of the meter periodicities when the highly repetitive stimuli were presented first, indicating a carry-over effect from the directly preceding condition, which offered maximal prior short-term context in terms of pattern repetition^[35]^.

### Periodization of rhythmic input in neural activity

Results revealed that the relative prominence of meter frequencies was enhanced in neural activity as compared to their relative prominence in the input. Notably, this enhancement was observed in all conditions, irrespective of the degree of pattern repetition in the input. Crucially, these neural responses cannot be explained by a mere tracking of prominent periodicities of the rhythmic input, as all stimuli were purposely designed to lack prominent temporal cues in the intermediate timescale corresponding to the beat layer (i.e., syncopated rhythms). This periodization of the rhythmic input observed in neural activity as captured with EEG thus arguably reflects processes related to meter perception, beyond lower-level sensory tracking of prominent features of the input.

Most importantly, the selective enhancement of meter frequencies was observed even in the non-repeating condition, which only offered a low degree of periodic recurrence in the input. This result suggests that periodic recurrence, even when limited only to faster timescales, is already sufficient to elicit significant neural emphasis of the meter. Together, these findings thus reveal processes through which the brain would leverage periodic recurrence restricted to a narrow timescale in the rhythmic input, to map a rich metric template covering wider timescales^[20,21,53]^. This multiscale temporal scaffolding would thus play a critical role in the neural periodization of rhythmic input that could be experienced as the meter^[9]^.

To some extent, these findings may be related to the phenomenon of subjective meter (or “tick tock” effect), which refers to perception of periodic accents in unaccented isochronous rhythms^[19,20,54]^. More specifically, it could be hypothesized that, in the no-pattern repetition condition, interpolation of the prominent periodic recurrences at the fastest inter-onset-interval timescale served as a basis to map an internal representation of a periodic structure at slower rates, thus yielding a set of nested periodic pulses. Previous studies^[55,56]^ on Western populations have shown a clear tendency towards tapping rates corresponding to groupings of 2, 4, and 8 notes (compared to groupings of 3, 5, 6, and 7 notes). Corroborating these observations, the current behavioral results showed a convergence towards duple meters.

However, the current study provides a nuanced differentiation from previous findings. Specifically, we found convergent tapping across participants at a rate corresponding to two times the fastest inter-onset intervals (Fig. 4). This tapping rate is thus overall faster, as compared to previous studies which mostly found tapping rates corresponding to four times these intervals in response to repeated rhythms at the same tempo^[8,10]^. The faster rate observed here thus likely reflects an attraction toward the most prominent periodic recurrence available in the sensory input, here located at the fastest timescale.

### Discrepancy between neural and behavioral responses

The behavioral measures showed less stable sensorimotor synchronization performances in non-musicians as compared to musicians, consistent with previous research^[2]^. This cross-group difference in the ability to tap a periodic beat could appear contradictory with the neural measures which did not show such a contrast between the two groups. Rather, the neural measures showed “periodization” of the input that was significantly influenced by pattern repetition and prior context, but not musical expertise.

Yet, these discrepancies between behavioral and neural measures actually align with the view of a default periodization of the rhythmic input in neural activity, which would contribute to meter perception irrespective of the musical expertise. Corroborating this view, previous research using intracerebral recordings in humans has provided evidence for such a periodization of rhythmic inputs at the earliest cortical stage of auditory processing^[26]^ (i.e., Heschl’s gyrus). These findings are also consistent with previous research showing periodization of syncopated rhythms in the EEG of human adults even when the attention is focused away from the rhythmic input^[43]^, as well as in the EEG of 5-6 months old human infants, despite relative immaturity of higher-level associative and sensorimotor cortices at this developmental stage^[41]^.

Taken together, these studies suggest that the capacity to move to a periodic beat does not only rely on a default periodized neural representation of the rhythmic input as produced in the primary auditory cortex. Rather, this ability would also require strong functional coupling between auditory and higher-level associative and sensorimotor cortices, allowing this periodized representation to guide movement timing, thus in line with the functional definition of meter in music^[57,58]^. In return, these processes would be reinforced through self-entrainment, especially in the context of syncopated rhythms. The critical role of such a functional coupling between these brain regions is further corroborated by the evidence that body movement can shape the internal representation of meter^[6,44,59]^.

In addition to the cross-group difference discussed above, we also found a cross-task difference. The positive effect of pattern repetition on the prominence of meter periodicities was found in the neural but not tapping response. This difference could be attributed to differences in task demands between behavioral and neural measurements involving, or not, explicitly instructed movement to the beat. Since the ability to move to the beat tends to be particularly developed in highly trained musicians due to lifelong music practice^[36,60]^, it could be hypothesized that musicians showed a ceiling effect in their behavioral responses, thus masking potential effects of pattern repetition. In non-musicians, in contrast, the overall weaker sensorimotor synchronization performances^[2]^, and the relatively low recurrence at slow supra-second timescales compared to previous similar studies^[8,10,43]^, may thus have proven too limited at enabling consistent motor entrainment to the beat, thus yielding a floor effect.

### Pattern repetition in music: capitalizing on implicit learning to foster meter perception

The current study shows that repetition of rhythmic patterns facilitates the internal representation of meter. This facilitation could stem from the capacity of the brain to subdivide the slow, supra-second periodicities provided by repetition of the rhythmic patterns into faster pulses, that is, a process of interpolation. Yet, this effect of pattern repetition is of a relatively small effect size. This relatively small effect could be explained by the fact that the slow timescales used here may have been already close to the limit beyond which periodic recurrence no longer facilitates meter perception^[4]^ (but see^[61]^ for extremely long timescale meter perception based on explicit theorization and training).

However, the human brain has proven to be remarkably proficient at recognizing and learning recurring patterns over time^[42,62,63]^. This phenomenon, referred to as implicit learning occurs on a short timescale already^[64]^, and does not require any specific training (see^[65]^ for a review of implicit learning in the context of music). Supporting this, effects of implicit learning were observed with auditory stimuli lasting up to 8.4 seconds^[42]^, thus comprising the duration of our “medium pattern repetition” condition (i.e. 4.8 s) used in the current study. The current results thus open to future research comparing neural and behavioral responses across a finer-grained range of durations of pattern repetition. For example, starting with approximately 2 s long patterns (as in^[8,10,26,41,43]^) could further clarify the role of implicit learning processes in fostering meter perception.

## Conclusion

Despite being a core component of music worldwide^[24,28–30]^, the repetition of musical patterns (melodic phrases, rhythmic figures, etc.) has often been overlooked in studies of rhythm processing. The current experiment aims to fill this gap by showing that the repetition of rhythmic patterns plays a critical role in fostering neural representations of temporal structures that can be experienced as the meter. Importantly, and beyond implications in music contexts, the obtained results reveal the capacity of the human brain to exploit slow periodicities available in the rhythmic input in the supra-second scale, to map internal representations of faster periodicities in sub-second timescales through interpolation. By highlighting the multiscale nature of temporal processes involved in rhythm processing, this study contributes to progress in our understanding of high-level perceptual processes allowing groups of individuals to coordinate interaction and communication in a multitude of fundamentally human joint actions ranging from music and dance performance to ritual, work, and play.

## Acknowledgements

The study is receiving financial support from the European Research Council (ERC Starting Grant Rhythm and Brains, ref. 801872). We would like to thank CATL, the technical support team of the Institute of Neurosciences at the Université Catholique de Louvain for providing the custom-built tapping box used in this experiment.

## Author Contributions

EC, TL and SN designed the study, and EC and SB collected and analyzed the data. EC, TL, RP and SN contributed to writing and editing the manuscript. All authors reviewed the manuscript.

## Code and data availability

Preprocessed and analyzed data are available on a public Zenodo repository (https://zenodo.org/records/13870035?preview=1). Additionally, anonymized raw data is available upon request.

The scripts used to conduct the analyses and figures are available on a public GitHub repository (https://github.com/Manu-RnB/pattern_repetition.git).

## Additional information

Correspondence concerning this article should be addressed to Emmanuel Coulon.

